# Cooperative population coding facilitates efficient sound source separability by adaptation to spatial statistics

**DOI:** 10.1101/536524

**Authors:** Helge Gleiss, Jörg Encke, Andrea Lingner, Todd R. Jennings, Sonja Brosel, Lars Kunz, Benedikt Grothe, Michael Pecka

**Affiliations:** Division of Neurobiology, Department of Biology II, Ludwig-Maximilians Universitaet Muenchen, Martinsried, Germany; Chair of Bio-Inspired Information Processing, Department of Electrical and Computer Engineering, Technical University of Munich, Garching, Germany

## Abstract

Our sensory environment changes constantly. Accordingly, neural systems continually adapt to the concurrent stimulus statistics to remain sensitive over a wide range of conditions. Such dynamic range adaptation (DRA) is assumed to increase both the effectiveness of the neuronal code and perceptual sensitivity. However, direct demonstrations of DRA-based efficient neuronal processing that also produces perceptional benefits are lacking. Here we investigated the impact of DRA on spatial coding in the rodent brain and the perception of human listeners. Naturalistic spatial stimulation with dynamically changing source locations elicited prominent DRA already on the initial spatial processing stage, the Lateral Superior Olive (LSO) of gerbils of either sex. Surprisingly, on the level of individual neurons, DRA diminished spatial tuning due to large response variability across trials. However, when considering single-trial population averages of multiple neurons, DRA enhanced the coding efficiency specifically for the concurrently most probable source locations. Intrinsic LSO population imaging of energy consumption combined with pharmacology revealed that a slow-acting LSO gain control mechanism distributes activity across a group of neurons during DRA, thereby enhancing population coding efficiency. Strikingly, such “efficient cooperative coding” also improved neuronal source separability specifically for the locations that were most likely to occur. These location-specific enhancements in neuronal coding were paralleled by human listeners exhibiting a selective improvement in spatial resolution. We conclude that, contrary to canonical models of sensory encoding, the primary motive of early spatial processing is efficiency optimization of neural populations for enhanced source separability in the concurrent environment.

**Author summary:** The renowned efficient coding hypothesis suggests that neural systems adapt their processing to the statistics of the environment to maximize information while minimizing the underlying energetic costs. It is further assumed that such neuronal adaptations also confer perceptual advantages. Yet direct demonstrations of adaptive mechanisms or strategies that result both in increased neuronal coding efficiency and perceptual benefits are lacking. Here we show that an auditory spatial processing circuit exploits slow-acting gain control to distribute activity across the neuronal population, thereby enhancing coding efficiency based on single-trial population averages. This population-efficiency maximization also results in improved neuronal spatial resolution for the concurrently most probable source locations, which was resembled in a focally improved spatial acuity of human listeners.

## Introduction

Our ability to distinguish individual objects in complex and dynamic environments is a fundamental brain function (1,2). Conversely, the functional requirements of sensory systems are shaped by the physical properties of the outside world: only if the neural sensitivity matches the current statistics of the sensory inputs, the coding of relevant stimulus features will be both informative and energetically efficient, and consequently evolutionary viable. Because realistic complex environments exhibit highly non-uniform occurrence probabilities of stimulus cues (3,4), sensory neurons adapt their action potential (“spike”) responses according to the probability of concurrent stimulus properties. This “dynamic range adaptation” (DRA) is thought to render neuronal firing maximally sensitive to changes in the stimulus range that is most likely to occur (Fig. 1) (5–7), while keeping activity rates low. Consequently, DRA to stimulus statistics is believed to reflect a neuronal adjustment to optimize stimulus encoding efficacy while simultaneously mediating improved perceptional resolution within the relevant cue range. However, direct demonstrations of DRA-based neuronal coding that causes both in increased neuronal efficiency and the resulting perceptual benefits are lacking.

**Figure 1:**
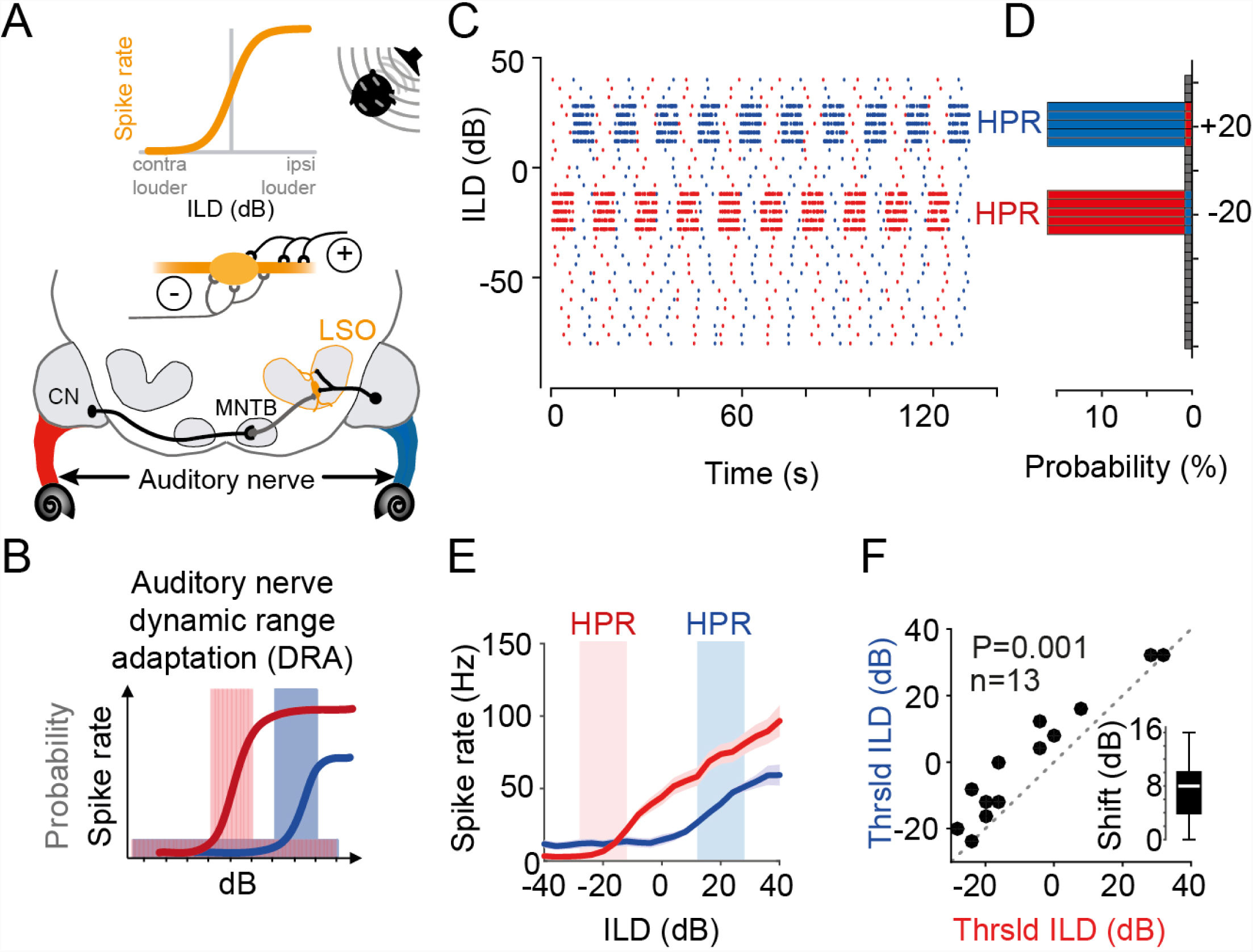
Statistic-dependent coding of spatial cues in LSO. A) Upper panel: LSO neurons respond with increasing higher spike rates to increasingly more ipsilateral sound source positions because these positions generate more “positive” ILDs (ILDs that favor the excitatory ear). Lower panel: Circuit diagram of the inputs to the LSO. LSO neurons receive excitatory input from the ipsilateral ear via the Cochlear Nucleus (CN) and inhibitory inputs from the contralateral ear via the Medial Nucleus of the Trapezoid Body (MNTB). B) Non-unimodal probability distributions of monaural stimulus intensity cause pronounced dynamic range adaptation already on the level of the auditory nerve fibers (ANFs) (16). Thus, for large ILDs, the ANFs in the left and right ear will adapt to different intensity levels (as indicated by color-coded ANFs in (a)). Colored shaded areas illustrate respective high-probability regions (HPRs) of stimulus value occurrence. See also (c) and (d). C) Illustration of the temporal sequence of ILDs in the two HPR conditions (centered on −20dB ILD, shown in red, or +20dB ILD, shown in blue) that were used to test the effect of complex stimulation on ILD coding. Note that the sequence changed for each of the 10 iterations of each HPR epoch. D) Probability histogram of the sequences shown in c). 80% of stimuli had ILDs centered on either −20dB ILD ±4dB (red) or +20dB ILD ±4dB (blue). E) Representative example of DRA in response to the change in HPR condition in a single LSO neuron. The ILD-response function for this neuron was substantially different between the 20dB ILD ±4dB (red) or +20dB ILD ±4dB (blue) condition. Given are mean response rates (solid lines) and S.E.M. (shaded area). F) Threshold ILDs (minimal ILD that significantly differed from baseline, see Methods) shift significantly by altering the HPR condition (P= 0.001, paired Wilcoxon signed rank test, n=13 neurons). Inset shows median shift (8dB, white bar) with IQR given by the box edges. Whiskers extend to overall data range. See also Figure S1.

In the auditory system, rapid (sub-second) DRA to stimulus statistics has been revealed on multiple processing levels from primary auditory cortex (8–11), to midbrain (12–15), and even brainstem (Fig. 1A). Specifically, DRA is prominently exhibited already by auditory nerve fibers (ANF) (16,17) (Fig. 1B), which consequently should affect the processing of all downstream centers, but might potentially be most crucial for spatial computations: To infer the location of a sound source, brainstem neurons of the Lateral Superior Olive (LSO) compare the difference in sound level at the two ears (interaural level difference, ILD) that is generated by a location-specific sound-attenuating effect of the head. LSO neurons respond according to the relative strength of excitatory and inhibitory inputs from the ipsi- and contralateral ear, respectively (Fig. 1A). The ensuing sigmoidal ILD-response functions (average action potential rate as a function of ILD) are regarded to represent the neuronal basis of auditory space encoding based on intensity difference cues (18) (Fig. 1A, top; in addition to timing-cues not dealt with in the present study). Specifically, individual source locations are thought to be mapped onto a specific spiking activity pattern of precisely tuned neurons or neuronal populations (19–22). Yet the nature of this spatial code and its read-out is still a matter of debate (23–26). Historically, studies have argued in favor of a labeled-line coding strategy of auditory space (or a mix of strategies), where small differences in the average spike rate tuning of individual, identifiable neurons or sub-population contributes to sound source localization (27–29). Yet, the majority of recent studies have concluded that sound source locations are initially encoded by the specific relative spike rate of two oppositely tuned hemispheric populations of spatially sensitive neurons (for review see (18)). This “two-channel hemispheric coding strategy” is motivated by the fact that the vast majority of neurons in each brainstem hemisphere is broadly tuned to contralateral sound sources, thus providing similar information about sound source locations (reviewed in (30)). The reasons for such apparently redundant and hence inefficient coding of space are, however, unknown. In either case, conclusions about spatial coding were derived from examining average neuronal firing rates in response to multiple repetitions of a stimulus set with uniform probability distributions of spatial cues (e.g. each ILD was equally likely to occur). Consequently, these traditional approaches neglected that under more natural conditions, DRA (of ANF or later stages) might crucially alter the nature of the neuronal code and/or its perceptual consequences:

First, the fact that sound source positions far to the left or right will result in distinctly different sound levels at the two ears (i.e. a large ILD) consequently should evoke DRA to different (monaural) stimulus levels for the ANFs in the left and right ear. Yet it remains to be tested how such differential monaural DRA impacts the detection and representation of ILDs in the LSO. Interestingly, we previously identified an activity-dependent LSO gain control mechanism (31) that might additionally influence the response to ILDs based on stimulation history (32). It follows that already the extraction of ILDs and consequentially the primary representation of auditory space might not be as rigid as traditionally assumed.

Second, the auditory pathway – like all sensory systems – must detect and code the relevant stimulus properties from only a single stimulus occurrence. The nature of response distributions to such a single instance of e.g. an ILD in a neuron population might be very different compared to the response distribution of a single neuron to this ILD averaged across trials, and consequently could result in different coding regimes. Since the extent of DRA can vary considerably across cells (10,16), such differences in response distributions might be particularly evident in complex acoustic environments.

Third, neuronal adaptations such as DRA to increase computational efficiency supposedly also entail a behavioral improvement within the concurrent environmental conditions. Hence, important insight into how the brain encodes auditory space under naturalistic conditions might be gained by investigating the perceptional impact of DRA (22). Yet while perceptional changes due to neuronal adaptation to stimulus statistics have been reported (15,33), demonstrations of task-specific perceptional benefits that are linked to neuronal efficiency and the underlying processing are missing.

To answer these questions, we studied the effects of naturalistic, i.e. spatially complex, stimulation on ILD processing in the LSO of gerbils and on the perception of human listeners. We extended a well-established monaural stimulus paradigm for studying DRA (10,12,16,17) and present rapidly changing ILDs that switched periodically between favoring either the left or right azimuthal space (Fig. 1C,D). In response to these spatially dynamic stimuli, we observed prominent DRA in LSO neurons, which demonstrates a lack of absolute encoding of space by average neuronal firing rate. Surprisingly, DRA in single neurons resulted in large response variability to a given ILD across trials. However, we find that when considering single-instance population coding, DRA maximized the efficiency of neuronal separability for specifically those ILDs that were most likely to occur in the concurrent statistical environment (high probability region, HPR, Fig. 1C,D). These stimulus-specific enhancements in neuronal coding were paralleled by human listeners exhibiting a selective improvement in just noticeable differences for ILDs within the respective HPR. Intrinsic LSO population imaging of energy consumption and a simple LSO model further explained that a slow-acting gain control mechanism enhances the population efficiency by distributing activity across a group of neurons during DRA. We conclude that, already on the primary detector level, the processing of ILDs is not tuned towards an absolute representation of space but optimizes efficient sound source separation in the concurrent acoustic environment by population coding of ILDs.

## Results

### LSO neurons exhibit DRA to spatial statistics

To explore the role of DRA on spatial coding in complex environments, we designed a stimulus paradigm with constantly varying ILDs in the context of two related but statistically distinct listening conditions. We used continuous broadband noise (identical on the two ears) and changed the ILD every 50ms, with ILD values drawn from one of two non-uniform distributions. The two distributions covered an identical range of ILDs but favored predominately (80% of time) either the ipsi- or contralateral ear (ILDs of +20±8dB and −20±8dB, named the +20dB HPR and −20dB HPR respectively, Fig. 1C, and Methods). This way, we simulated dynamic spatial environments with dominant sound sources located either left or right of midline (Fig. 1D). The two conditions switched periodically (Fig. 1C, one run consisted of 19 switches every ~6s, the sequence of ILDs was different for each switch but identical across repetitions, 3 runs were recorded for each cell). To assess to what extent changes in spatial statistics alter the neuronal detection and encoding of ILDs, we first carried out extracellular recordings from single neurons in the LSO of anesthetized gerbils while presenting the stimuli via calibrated earphones (see Methods).

Following previous studies of DRA (12,14–16), we first assessed the average neuronal spike rates (calculated across occurrences of each ILD) separately for −20dB to the +20dB HPR conditions. We observed that the resulting ILD-response functions differed between the two conditions (a single neuron example is shown in Fig. 1E). Specifically, a clear shift of the ILD-spike rate functions was observable that entailed a change in the average spike rate in the respective HPRs (red and blue areas in Fig. 1E and throughout). To quantify these shifts, which appeared highly reminiscent of DRA to accommodate the change in the range of overrepresented ILDs, we computed the minimal ILD that triggered significant spiking (“threshold ILD”, see Methods) in the respective condition for each neuron. Threshold ILDs significantly increased when switching from the −20dB to the +20dB HPR (Fig. 1F, n=13 neurons, P=0.001, paired Wilcoxon signed rank test). For the population, the median shift in threshold ILD between the two conditions was 8dB (interquartile range [IQR] 16dB; Fig. 1F inset). To further characterize the extent and specificity of the DRA, we also generated two additional “un-natural” ILD distributions. These additional conditions confirmed that the observed shifts in threshold ILDs were dependent on the concurrent binaural statistics (Fig. S1, n=18 neurons and n=19 neurons). This presence of DRA-related shifts in ILD-functions in the LSO directly demonstrates a lack of absolute encoding of sound source locations by the average neuronal firing rate already on the level of cue detection.

### DRA optimizes single observation population coding

So far, we followed previous studies of DRA in the auditory system (12,14–16) and evaluated the spatial sensitivity of individual LSO neurons by its average spike rate given the repeated presentation of each ILD. However, in reality, processing must be able to compute the location of a sound source from observation of a single instance of the stimulus. Therefore, we next focused on the direct response by each neuron to each occurrence of a particular ILD. Examining individual spike counts for 75 recurrent instances of +20dB and −20dB ILD in the respective HPR condition revealed two interesting findings (Fig. 2A):

**Figure 2:**
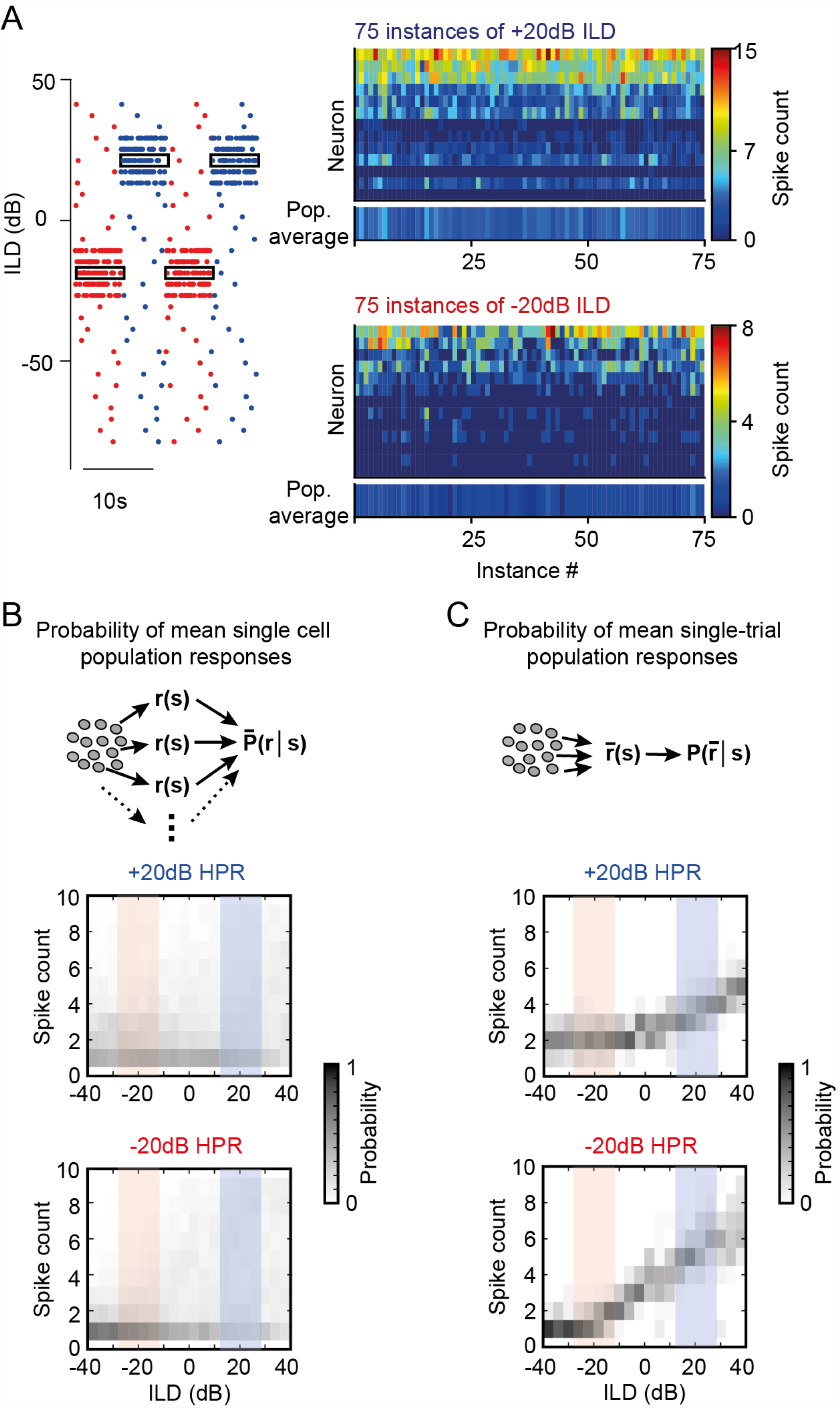
Statistic-dependent DRA results in sparse and selective ILD coding for single-instance population averages. While response probabilities were low overall, the variability in spike counts to repetitive instances of the same ILD in individual neurons was high. Shown are spike counts of all neurons in response to 75 instances of −20dB ILD (lower right-hand panel) or +20dB ILD (upper right-hand panel). Bottom line in each panel shows mean responses for each ILD instance. and C) Mean population spike-count probability density functions of all neurons were constructed based on the pooled response of all single neuron responses (B) and population mean responses at each instance of ILD occurrence (C). Only the later resulted in informative ILD tuning (see also Fig. 3 and Fig. S2).

First, responses of most LSO neurons within the concurrent HPR were very sparse (median spike count and IQR: −20dB HPR: 0.92 and 0.46 spikes; +20dB HPR: 3.15 and 0.85 spikes). Second, a high response variability, as indicated by the large IQR, was observable for all ILDs: spike counts varied considerably between repeated instances of the same ILD (trial-wise median Pearson’s correlation coefficient and IQR: 0.62 and 0.26, Fig. S2) and spike-triggered average analysis showed no systematic relationship between ILD sequences and their likelihood to trigger a spike (Fig. S2). Crucially, this lack of consistent responses to ILDs with naturalistic probability statistics resulted in a very limited modulation of the average spiking probabilities in either HPR condition. That is, the probability to observe a particular mean average spike count across the sample population was very similar for all ILDs (Fig. 2B).

In contrast, however, more specific ILD population tuning emerged from our data set when considering the population response for a single occurrence of a particular ILD (i.e. averaging across a column in Fig. 2A; Fig. 2C, compare also bottom lines in right-hand panels of Fig. 2A; note that neurons were recorded sequentially). To determine how these different population tuning of mean single cell population responses and single instance population responses impact the decoding accuracy of ILDs, we performed a Maximum Likelihood Estimation (MLE, see Methods) for both methods. In short, MLE approximates which ILD is most likely to have occurred given the observation of a particular spike count. The estimates differed in two important ways when either considering the mean single cell population responses (MLE(mean)) or the single instance population response (MLE(pop)) (Fig. 3A,B): First, the estimated deviation from the true ILD values were much larger for the MLE(mean) (minimal deviations: −20dB HPR: 14.3 dB @ −24dB; +20dB HPR: 16.8 dB @ −16dB) compared to MLE(pop) (minimal deviations: −20dB HPR: 4.7 dB @ −20dB; +20dB HPR: 10.1 dB @ 0dB); i.e. the accuracy of MLE(pop) was higher. Secondly, the generally assumed advantageous effect of DRA, i.e. that DRA explicitly enhances the coding in the respective HPR was evident for MLE(pop), but not MLE(mean) (Fig. 3A,B; improvement of 9dB and 0dB, respectively).

**Figure 3:**
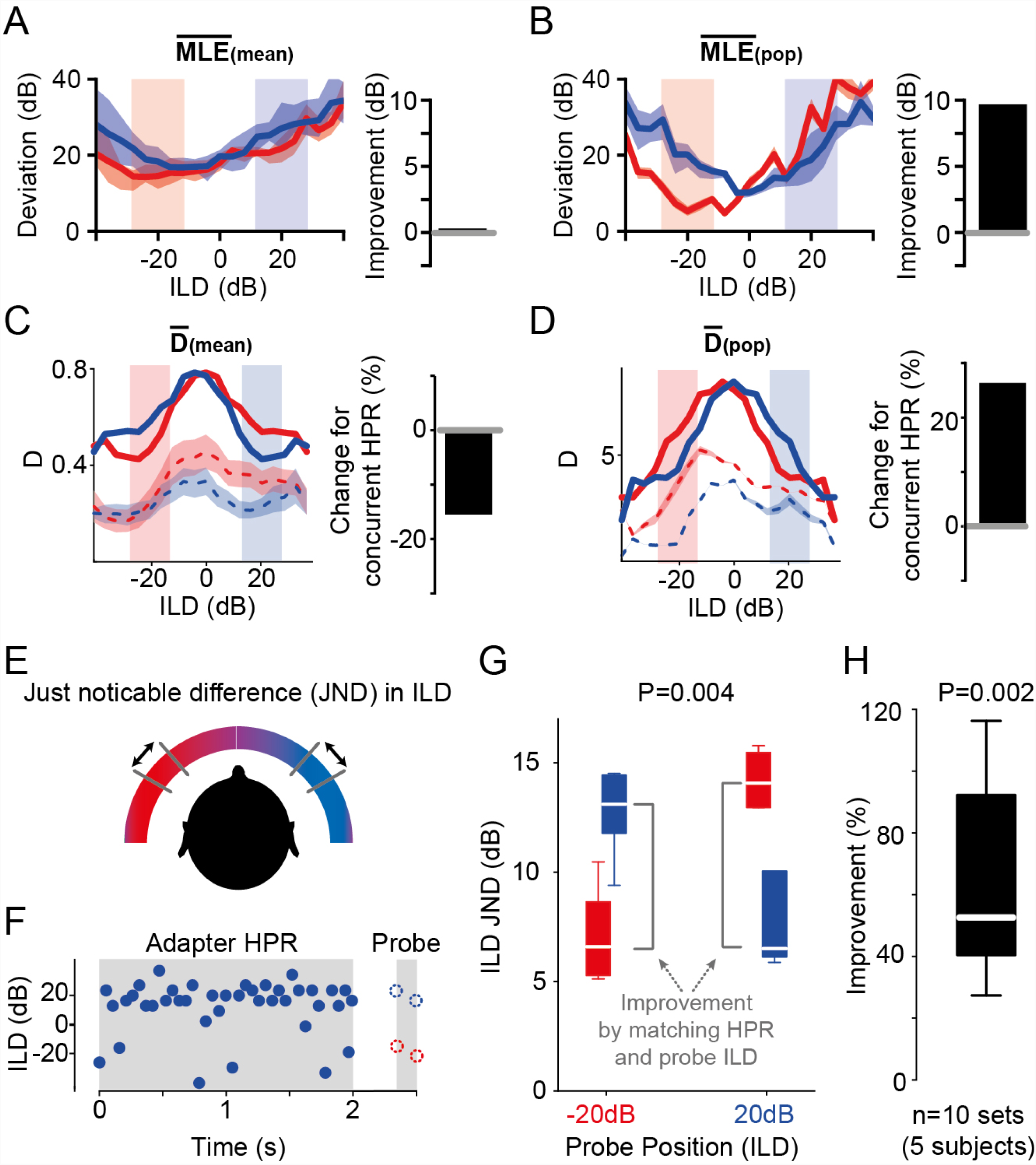
Single-instance population averaging predicts human spatial perception. A) Average MLE(mean) exhibited no apparent advantage of DRA for decoding ILDs (improvement = 0%, right-hand panel), as the tuning function obtained from the −20dB HPR condition was overall better in estimating the ILDs compared to those obtained from the +20dB HPR condition. Solid lines show the mean deviations and shaded area represents S.E.M.. B) Average MLE(pop) showed a 9dB improvement (calculated as the difference between the red and blue functions within the HPRs) for decoding ILDs within the concurrent HPR. Conventions as in (A). C) Average D(mean) across all cells (n=13) decreased for the concurrent HPR, suggesting a relative worsening of separability of ILDs of by −15.5% (right hand panel). Solid lines represent D including the hypothetical second hemispheric response. Mean data from single hemisphere are given by dashed lines (shaded area represents S.E.M.). D) Average D(pop) of the recorded LSO neurons (n=13) increased within the concurrent HPR by 27.1% (right-hand panel). Solid lines represent D(pop) including the hypothetical second hemispheric response. Mean data from single hemisphere are given by dashed lines (shaded area represents S.E.M.). E) and F) The effect of DRA to HPRs on ILD JND of human subjects were measured with stimuli presented over headphones. Presentation of an adapter sound consisting of a 2s snippet of one of the two HPR stimuli was followed by two noise probes (50ms each) for JND measurement. The ILDs of the probe tones were centered on −20dB ILD or +20dB ILD, and hence either matched or miss-matched the HPR of the preceding adapter. G) Single-subject example of the influence of the HPR on ILD JND. Co-location (i.e. matching) of the adapter HPR and probe sound position led to significant improvement of the ILD JND (P=0.004, Friedman test, n=12 trials each). H) The average improvement of co-location by adapter HPR and probe tone ILD across listeners was 52.6% (P=0.002, unpaired Wilcoxon signed rank test, n=10 sets from 5 subjects, conventions as in inset in Fig. 1F).

Such relative enhancement in the neuronal precision of ILD estimation implies – but does not confirm – a relative improvement in the ability to resolve nearby sound locations. To quantify the impact of the differences in population tuning (Fig. 2B,C) on resolution directly, we next determined the informational content of each neurons’ response towards the separability of ILDs. We followed previous studies on ILD coding (15,34) and calculated the standard separation (35) (“D”), which quantifies the separability of adjacent ILDs based on the ratio of slope steepness and response variability. We first calculated D(mean), i.e. using the mean of average single neuron tunings. Since the two spatial conditions that we presented were mirror-symmetric to each other, the two LSOs on each side of the brain would provide complementary spatial information for each HPR condition towards D. We therefore summed the D-ILD functions of each condition with the mirror-image of the function of the other condition (Fig. 3C, dashed lines indicate single-hemisphere data, solid line represents sum of both hemispheres). In remarkable contrast to previous midbrain studies (12,15), we found that the average neuronal separability was not enhanced by the DRA, but actually considerably lower within the respective HPRs (Fig. 3C; change for concurrent HPR: −15 %, compare red and blue lines in respective HPRs in left panel). Thus, D(mean) would predict a worsening of ILD resolution by the observed DRA to spatial stimulus statistics. In contrast, when D is calculated based on the mean spike count of all neurons to each instance of an ILD (D(pop)), high specificity of separability for the ILDs of the concurrent HPR becomes evident (Fig. 3D, change for concurrent HPR: +27 %).

### Human spatial resolution improves specifically for HPR ILDs

To test whether the increased performance as predicted by analyzing MLE(pop) and D(pop) also results in an improved ability to resolve sound source locations, we performed a spatial separability test with human listeners via calibrated headphones (Fig. 3E). The subjects (N=5) were presented with a 2s long snippet of the same stimulus used in the electrophysiological experiments, taken alternatively from the +20dB and −20dB ILD HPR condition (only +20 dB is illustrated in Fig. 3f). Shortly after (0.35s) this adapting period, the listeners were presented with two probe ILDs (each consisting of 50ms broadband noise, spaced apart by 100ms) and were asked to indicate which of the two was perceived more lateralized. Using an adaptive tracking paradigm (see Methods), the difference in ILD between the two probe ILDs was systematically reduced to determine the just noticeable difference (JND) in ILD for each subject. The probe ILDs were centered either on +20db ILD or −20dB ILD (Fig. 3F), to allow deciphering the influence of matching and miss-matching the adapter HPRs with the probe ILD. In agreement with the prediction made on the basis of D(pop), we observed a significant improvement in JND when probe center ILDs matched the HPR of the adapter (single subject example in Fig. 3G: −20db adapter and −20dB probe, red, or +20dB adapter and +20dB probe, blue; P=0.004, Friedman test). On average, JNDs of the five listeners improved by 52.6% (Fig. 3H, IQR: 51.8%, P=0.002, Wilcoxon signed rank test). Thus, human JND performance is in close agreement with the neuronally derived MLE(pop) and D(pop), suggesting that DRA crucially affects both population coding and perception of ILDs in complex acoustic environments.

### Slow gain control maximizes efficiency

How could single instance population responses in the LSO be optimized for separability of ILDs in the concurrent HPR? Moreover, what effect might link high response variability of single neurons and highly informative population ILD coding? To gain insight into potential underlying mechanisms, we first analyzed the time course of DRA in LSO neurons. In accordance with DRA studies using similar stimulus statistics in other centers of the auditory system(12,14,16), we observed an exponential time course of rate adaptation (Fig. 4A). Yet in contrast to all previous reports, we found that adaptation kinetics were best described not by a single, but by two time constants (Fig. 4B). Addition of the second time constant resulted in lower root mean squared errors of fits (rmse, median rmse_(double)_=0.15, median rmse_(single)_=0.26, P=0.001, Wilcoxon signed rank test; Fig. 4C) and double exponential fitting was significantly superior to single exponential fits even after compensating for the unspecific benefit of an additional fitting parameter (Fig. 5d, median adjusted R^2^_(double)_=0.45, median adjusted R^2^_(single)_ =0.025, P=0.001, Wilcoxon signed rank test). The shorter (first) time constants of the fit (Fig. 4B; median tau = 222.8ms, IQR: 1.465s) were similar to the kinetics reported for the auditory nerve (17), suggesting that monaural DRA upstream of the LSO contributed substantially to the ILD adaptations in the LSO. The second time constants were considerably slower and in the range of a few seconds (Fig. 4B; median tau = 2.2s, IQR: 7.7s).

**Figure 4:**
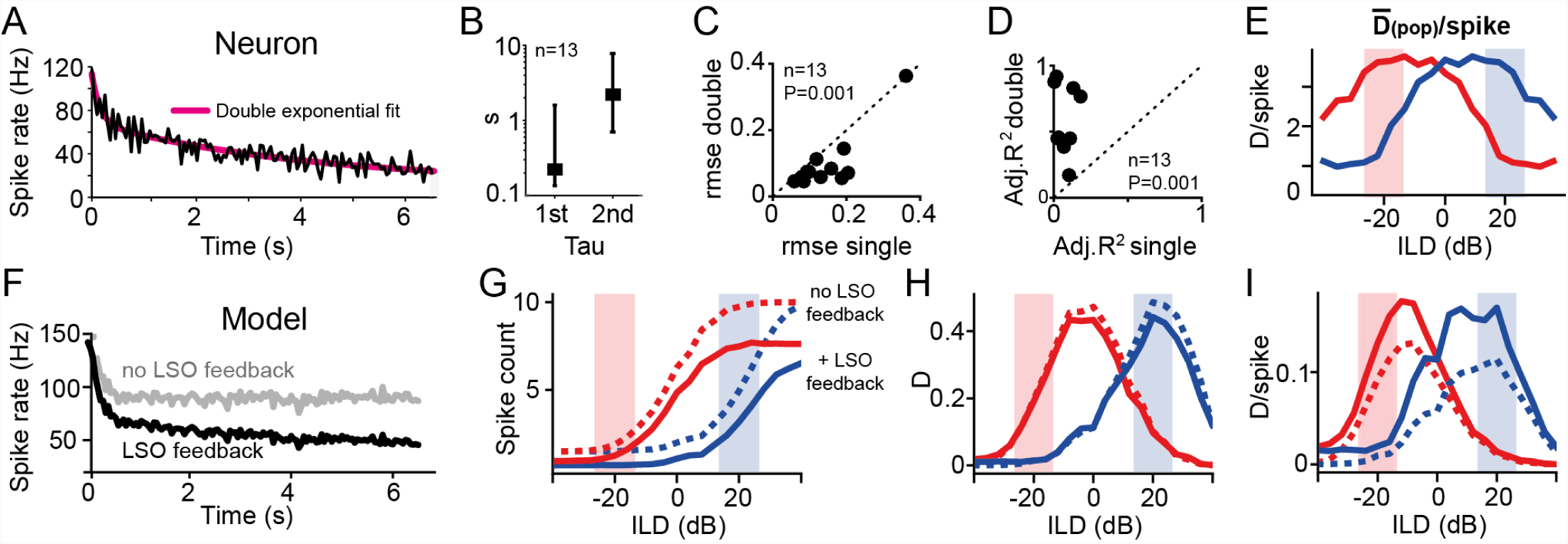
Slow rate adaptation increases population efficiency. A) Evolution of rate adaption is best explained by a double-exponential process. Shown is a single neuron example (black) and double-exponential fit (magenta). Beside a rapid rate adaption at the beginning of an epoch, a second, slow time constant of was also present in gerbil LSO. B) Average fast and slow time constants of DRA in the LSO from double exponential fitting: median tau(1^st^) = 0.22 s, IQR: 1.465s; median tau (2^nd^) = 2.2s, IQR: 7.1s. C) and D) Inclusion of a second time constant was superior to single time constant fitting of rate adaptation: the root-mean squared errors (rmses) of fits decreased ((C); P=0.001, paired Student t-test, n=13 neurons) and adjusted R^2^s increased ((D); P=0.001, paired Student t-test, n=13 neurons). E) Efficiency of responses was measured by calculating D(pop)/spike, displaying high specificity for ILDs in the concurrent HPR. F) A simple subtraction model of the LSO and DRA of its inputs can replicate the electrophysiological results only when including a binaural gain control stage. Right panel shows a diagram of LSO model including negative feedback after binaural integration. Left panel shows adaptation time course of the model (as for the neuron in (A)). Gray and black traces represent results excluding and including binaural negative feedback, respectively. G), H), I) The model was able to qualitatively reproduce both the HRP-specific shifting effect of DRA in the LSO (G) and the HRP specificity of D (H) and D/spike (I). Model responses without and with slow negative feedback stage after binaural processing are shown by dashed and solid lines, respectively. Crucially, the presence of negative feedback resulted in stronger rate adaptation, which did not affect D, but substantially increased efficiency by 55% (mean improvement within the two HPR regions).

**Figure 5:**
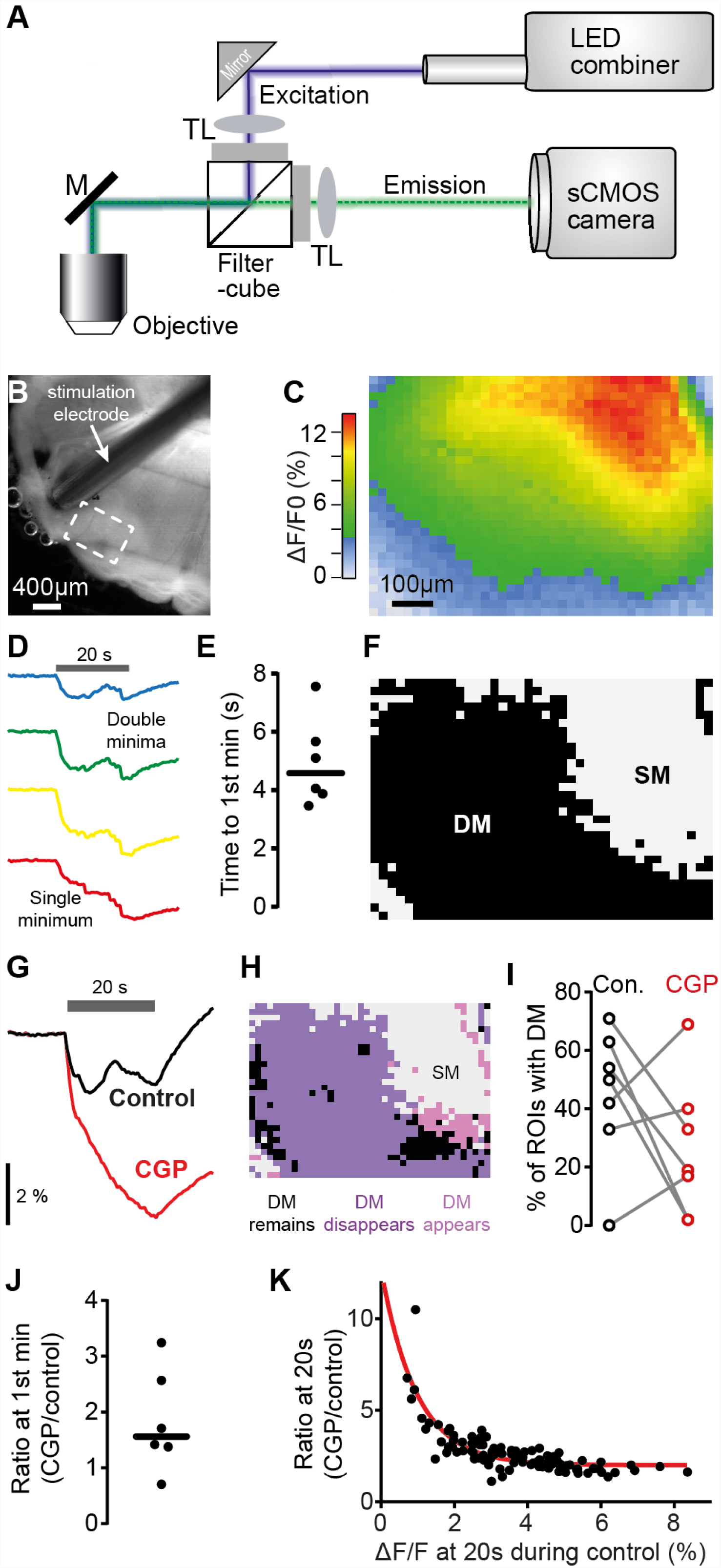
GABA-B signaling regulates NADH consumption in LSO in activity-dependent manner. A) Schematic view of the optical path for intrinsic fluorescence imaging of metabolic activity in the LSO. M: mirror; TL: tube lens. B) Bright-field image of brainstem (oblique illumination with IR-LED) Dashed rectangular denotes imaged area shown in (C). C) Heat map illustrating spatial distribution of maximal relative decrease of NADH autofluorescence in the LSO in response to 20 s fiber stimulation at 200 Hz. D) Exemplary traces for the temporal evolution of changes of NADH levels (colors correspond to respective regions in (C). Gray horizontal bar denotes duration of electrical stimulation. While only a single minimum (SM) was present in the bottom trace (red, corresponding to large NADH decrease), the top three traces with less overall NADH decrease exhibited double minima (DM). E) Mean time from stimulation start to 1^st^ minimum calculated for each slice from those ROIs that exhibited DMs. Population median: 4.58 s. F) Distribution of SMs and DMs in the recording shown in (C). G) Exemplary CGP-induced change in NADH fluorescence of a single ROI. H) Spatial distribution of CGP-induced changes of NADH response types (SM / DM) for the recording shown in (C). I) CGP-induced changes in the fraction of ROIs showing DMs (n = 6 slices). J) The average CGP-to-control ratio of NADH levels at mean time point of 1st minima (independent of presence of 1st minimum in the respective ROI) was 1.6. K) CGP-to-control ratio of NADH levels as function of NADH level changes in the control measurement (both measured 20s after onset, in 100 neighboring ROIs in (C); red line corresponds to exponential fit (tau=0.97%; Spearman correlation, r = 0.8176, P <0.0001).

Such slow rate adaptation is consistent with previous reports of negative feedback loops in LSO neurons: the inhibitory transmitter *gamma*-aminobutyric acid (GABA) is released in an activity-dependent manner into the extracellular matrix and thereby suppresses pre-synaptic inputs in the vicinity via slow-acting GABA-B receptors (31). Hence, DRA in the LSO might be considerably influenced by a slow binaural gain control for ILD coding during naturalistic stimulation. It had been suggested that such slow negative feedback serve to increase the efficiency of population coding (36,37): Because spiking is energetically costly, the efficiency of a neuronal representation depends on the informational content of a spiking response relative to the number of spikes that conveyed this information (38,39). To quantify neuronal population efficiency directly, we calculated the average D transmitted per spike for the instantaneous hemispheric average (D(pop)/spike). This analysis revealed a hemispheric specificity of response efficiency for the concurrent spatial conditions (Fig. 4E). Thus, the slow gain control mechanism associated with the second time constant of DRA that we found might serve to maximize the efficiency of neuronal processing within the range of most likely occurring ILDs. To investigate this potential role of slow gain control on ILD coding in more detail, we generated a simple model of the LSO based on existing models of DRA. Specifically, we extended an existing auditory nerve model that included both threshold and gain adaptation (17) by adding a binaural subtraction stage to reflect LSO processing (Fig. S3). As expected, this simple model exhibited clear DRA in response to the binaural HPR stimuli (Fig. 4G, dotted lines). However, since this version of the model lacked a binaural gain control stage, it captured only the fast time course of rate adaptation and quickly reached a steady state spike rate (<1s, Fig. 4F, gray trace). To account for the second, slow adaptation component in the neuronal data, we included an additional slow negative feedback stage after binaural comparison in the model (Fig. S3). This modification resulted in a close match in the dynamics of rate adaptation between model and LSO neurons (Fig. 4F, black trace) and led to lowered overall spike counts during DRA (Fig. 4G, solid lines). This effect of slow gain control had little effect on the overall amount of spatial information (Fig. 4H), but specifically increased D/spike of model responses within the concurrent HPR (Fig. 4I, compare dotted and solid lines). These modeling results thus suggest that the main function of slow gain control in the LSO is the optimization of coding efficiency (i.e. separability per unit of neuronal activity).

### Intrinsic imaging reveals energetic benefits of slow GABAergic gain control

To directly test the model prediction that slow feedback signaling may minimize energy expenditure in the LSO, we took advantage of the intrinsic autofluorescence of NADH as key metabolic intermediate (40,41). We monitored the relative change in NADH levels with high spatial resolution in LSO brain slices (21µm x 23µm per region of interest, “ROIs”, 1200 ROIs per field of view; Fig. 5A,B, see Methods). Using 20 second long fiber stimulation at 200 Hz, we determined the spatial distribution of energy consumption in the LSO (6 brain slices). As expected, large parts of the imaged LSO area displayed a monotonic increase in energy consumption with a single minimum (SM) in response to the 20 second long stimulation (Fig. 5C, red region, Fig. 5D, lowest trace). However, we also frequently observed areas in which energy consumption declined after a few seconds of stimulation, before ultimately increasing again (“Double minima, “DM”, Fig. 5D, and Fig. 5F). This non-monotonic progression of energy consumption combined with its apparent slow time course (4.58s; IQR: 2.36s, Fig. 5E) is highly suggestive of the known GABA-B receptor mediated, activity-dependent gain control mechanism. Accordingly, application of the specific antagonist CGP 55845 hydrochloride (CGP, 10µM) to the bath revealed that DMs largely disappeared during blockade of GABA-B signaling, resulting in considerable larger energy consumption (Fig. 5G). In accordance with the assumed gain control function of GABA, on the population level (i.e. across all ROIs per slice), CGP had differential effects on the prevalence of observed DMs. A spatial diversity in the effect of blocking GABA-B mediated inhibition was clearly observable within individual brain slices (Fig. 5H). Specifically, DMs were either more or less likely to appear during CGP dependent on the fraction of DMs during control (Fig. 5I).

Application of CGP also had a striking effect on the overall energy consumption in the LSO: Across the entire imaged area, the block of GABA-B signaling on average almost doubled the energy consumption (median CGP-control ratio: 1.6, IQR: 1.5; n= 6 slices, Fig. 5J). Moreover, similar to the history-dependency observed for the DMs, the magnitude of change in the energy consumption during CGP application was highly correlated with the prior activity level during control conditions (Spearman correlation, P<0.0001, Fig. 5K), providing further corroboration for the activity-dependency of the gain control mechanism.Together, these data strongly suggest that the spatially variable, slow gain control mediated by GABA-B in the LSO serves for the efficient population coding of ILDs.

## Discussion

Our findings advocate a novel concept for the neuronal coding of auditory space that significantly differs from the canonically assumed mapping of absolute locations (42). We observed that LSO neurons strongly adapted their ILD-rate functions in response to changes in the spatial statistics. Consequently, auditory spatial representation is dynamic and devoid of absolute mapping of sound source locations already on the detector level. We further discovered that the average rate tuning of single LSO neurons conveys little spatial information during naturalistic stimulation due to high response variability. However, if responses to individual instances of an ILD were averaged across neurons, DRA optimized the efficiency of responses, which resulted in improved separation of ILDs within the concurrent HPRs. Correspondingly, human listeners showed evidence of a focal improvement in ILD resolution specifically for HPR ILDs. Importantly, this study is – to our knowledge – the first to demonstrate stimulus-specific benefits by DRA both for the efficiency of neuronal coding as well as human perception. Finally, a simple LSO model and intrinsic energy imaging explained that the efficiency of the enhancement in spatial separability is facilitated by a slow gain control mechanism involving GABAergic signaling downstream to binaural integration.

The canonical concept of spatial coding assumes that specific average response rates of sensory neurons are mapped onto a particular physical cue to allow for a faithful encoding of the corresponding source location (19–21). By introducing a more naturalistic likelihood-distribution of spatial statistics, we determined a prominent role of DRA for binaural processing that refutes the idea of an absolute representation of space already on the detector level and instead promotes a relative coding of sound source positions. Moreover, our evaluation of the impact of DRA on neuronal information suggests that the basic principle of LSO spatial coding is the preservation of ecologically relevant coding efficiency by providing high separability of nearby sound sources within the statistically predominant range of ILDs (5,38). These findings corroborate the generality of these perceptual effects of stimulus history independent of the specific binaural cue(42), and provide an mechanistic explanation on the detector level that had so far been linked to secondary processing at higher stages (13,15,43–45).

Because the LSO represents the initial binaural stage of ILD detection, our findings stand out for two more reasons: (1) Adaptive processing at the spatial cue detector should result in absolute localization errors due to a missing reference frame. This notion is supported by reports of human listeners producing significant absolute localization errors when presented with biased spatial statistics (33,42,44,46). (2) While adaptation with the purpose to preserve a large dynamic coding range within the predominant stimulus range can be found across sensory systems (7), we showed that in the auditory spatial system, DRA is likely to be inherited to a large degree by adaptation to intensity statistics in the monaural LSO inputs (e.g. the auditory nerves from each ear). We furthermore show that the major computational modification after binaural integration serves to optimize the efficiency of coding for the concurrent spatial statistics. Such processing of naturalistic statistics to maximize the efficiency of information transmission (by redundancy reduction) had so far been associated with midbrain and cortex, i.e. processing that is secondary to the initial detection of the respective feature (47–51). In contrast, ILD detection and efficiency optimization are realized concurrently by the LSO (and subsequent negative feedback, see below). Interestingly, adaptation to binaural statistics to optimize spatial sensitivity has also been described for the detector neurons of the second important binaural cue, the interaural time differences (28,52). However, in contrast to the short-term changes of the LSO, these adaptations take place over days during maturation and entail long-term morphological changes.

Different to prior studies on adaptation to spatial statistics in the midbrain (12,15), MLE(mean) declined within the concurrent HPR due to the high response variability of individual neurons. An informational gain was only revealed by applying a single observation population coding concept in the form of MLE(pop). In this regard, our data provide physiological support for the framework of cooperative population decoding (36), which had been developed to explain the apparent noisiness of cortical processing. Specifically, the framework suggested that recurrent inhibition with slow time constant can be utilized to maximize the efficiency of an average population code to the expense of increased response variability of individual neurons. Congruent with such a coding regime, individual LSO neurons responded sparsely (intermitted and with few spikes) and therefore decreased the redundancy of firing in the population for a given ILD. A potential limitation for such an interpretation of our data is that the population of neurons was not recorded at the same time (also due to methodological limitations for brainstem recordings of highly stimulus-time-locked responses). However, our *in vitro* recording of large population of LSO neurons conclusively supports the single neuron data and moreover, it is known that spiking in auditory brainstem nuclei occurs independently (53). Accordingly, population analyses of single neuron recordings are assumed a valid approximation and thus commonly performed (12,13,15).

In conclusion, our findings suggest a new concept for auditory spatial coding: the detection and processing of spatial cues, particularly ILDs, is not geared for an absolute representation of space, but optimizes efficient sound source separation in a given stimulus context by sparse population coding.

## Materials and Methods

### Electrophysiology

*In vivo* extracellular single cell recordings were made from the LSO of young adult (postnatal age >80 days, n = 7 animals) Mongolian gerbils (*Meriones unguiculatus*) of both sexes. The experiments were approved by the German animal welfare act (District Government of Upper Bavaria, reference number: 55.2-1-54-2531-105-10).

To anesthetize the animals, a combination of ketamine (Ketavet, 100 mg/mL; Pfizer Inc., USA) and xylazine (Xylazin, 100 mg/mL; Sigma-Aldrich Chemie GmbH, Germany) was used. Physiological sodium chloride solution (NaCl, 0,9 %, B. Braun Medicare GmbH, Germany) was mixed with 20 % of ketamine and 2 % of xylazine. After weighting the animals they were anesthetized with an intraperitoneal injection (0.5 ml per 100 g body weight) of this anesthetic. After initial injection, the anesthetic was continuously provided by an automatic pump (801 Syringe Pump, Univentor High Precision Instruments Ltd., Spain) at a rate of 1.6 to 2.8 μl per minute depending on body weight and state of anesthesia. The anesthetic stage was periodically tested with the hind leg reflex. Constant body temperature of 37 °C was ensured and checked by a thermostatically controlled heating pad the animals were placed on (Harvard Homeothermic Blanket Control Unit Model #50-7129, Harvard Apparatus Inc., USA). In order to ensure a sealed placement of the headphones on the acoustic meatus the tragus was cut at two sides. The pericranium was anesthetized with Lidocaine (Xylocain Pumpspray dental, AstraZeneca GmbH, Germany).

A small cut of the skin was made across the rostro-caudal axis on the upper part of the skull and a craniotomy and a durotomy (ca. 1.5 x 2.5 mm) approx. 1800 mm lateral to the midline and 4500 mm caudal to the bregmoid axis was performed. Ringer solution was periodically applied to the opening to prevent damage of the brain surface due to dehydration. The animals’ body functions were monitored though various devices. The heart rate and breathing cycle was monitored optically and acoustically through a electrocardiogram. The animals’ blood oxygen was measured through a pulse oximetry monitor (LifeSense Tabletop Capnography and Pulse Oximetry Monitor, Nonin Medical Inc., USA). The animal was also typically provided carbogen through a custom-made mask. Recording sessions typically lasted between 10 to 12 hours. The recording site was marked by iontophoretic application of the enzyme horseradish peroxidase (HRP). Experiments were then finalized by euthanizing the animals without awakening by an intraperitoneal injection of 1 ml of 20 mg/ml pentobarbital in ringer solution. The animals were transcardially perfused with Ringer solution and 4 % paraformaldehyde (PFA) for approximate 30 minutes. The brain was carefully removed from the cranium and put into 4 % PFA at 4 °C upon further processing to determine the recording location. Only recordings from locations that were positively identified within the LSO were used for further data analysis.

Extracellular single-cell recordings were obtained using pulled glass micropipettes (1.5 mm O.D. x 0.86 mm I.D., GC150F-10, Harvard Apparatus Ltd., USA) filled with 1 M HRP in 1 M NaCl and a resistance of 7 to 10 MΩ (measured with Ωmega-Tip Z, World Precision Instruments Inc., USA). The electrode was mounted on a piezo drive (Inchworm controller 8200, Burleigh Products Group Inc., USA), which was connected to a motorized manipulator (Digimatic series 164 type 161, Mitutoyo Deutschland GmbH, Germany). The electrode signal was amplified (Electro 705, World Precision Instruments Inc. and Wide Band Amplifier, TOE 7607, Toellner GmbH, Germany) and fed to a computer via an A/D-converter (TDT RP2.1, System III, Tucker-Davis Technologies Inc., USA) where the signal was filtered. A notch filter was used to filter the 50 Hz electrical noise caused by the power line hum and a high-pass filter with 300 Hz and low-pass with 5 kHz bandpass filtered the signal in the RP2.1. Brainware (Jan Schnupp, University of Oxford UK, for Tucker-Davis Technologies Inc., USA) was used to visualize and analyze incoming spike trains. The spike times and raw traces were recorded and saved for subsequent analysis.

### Stimulus generation and presentation

All the stimuli were digitally generated using MatLab (The MathWorks Inc., USA) and fed into TDT hardware using Brainware. The stimuli were D/A-converted in a TDT Multi-Function Processor (TDT RX6, System III, Tucker Davis Technologies Inc., USA) and then attenuated with a TDT Programmable Attenuator (TDT PA5, System III, Tucker Davis Technologies Inc., USA). The analog signal was delivered to the head phones. To cover the a wide range of the LSO’s frequency spectrum, either Etymotic Research headphones ER-10B+ with ER 10D-T04 silicon ear tips (Etymotic Research, Inc., USA) or custom-build electrostatic headphones were used. The same silicon ear tips where fitted to either headphones to have a comparable seal to the animals ears. Custom-written calibration filters were used to achieve a flat spectrum over the entire range of the respective headphones. When spikes of single cells were identifiable, the characteristic frequency (CF) and absolute threshold were determined using a pure-tone stimulus having the same length as the search stimulus. For further characterization of a neuron, a baseline ILD function was obtained and a broad-band noise rate-level functions were recorded in response to 50ms bursts presented on the ipsilateral excitatory ear only. These recordings were used for determining a cell’s latency (see below). To measure DRA in LSO neurons, a bimodal HPR stimulus was created. The intensity of continuous broadband noise was drawn from a pseudo-randomized predefined distribution every 50 ms (see Fig. 1). The range of monaural intensities spread from 20 to 80 dB SPL in 2 dB steps, similar as used monaurally (16,17). The predefined distribution consisted of two HPRs intensity levels around center-intensities of 50 dB ±4 dB SPL and 70 dB±4 dB SPL, resulting in 5 values per HPR with a cumulative occurrence probability of 0.8. To generate ILDs, the stimulus intensities were mirrored at 60 dB for presentation on the other ear, resulting in HPR center regions of −20 dB ILD and +20 dB ILD, respectively. A single condition epoch was 6.55 seconds long and was repeated ten times with different pseudo-randomizations each time (but identical cumulative probabilities). A 131 seconds long stimulus was generated by alternating the two HPR conditions repeatedly, resulting in 10 HPR epochs per sweep for each condition.

### Neuronal data analysis

Recorded data files were analyzed offline using custom-made analysis in MatLab and Python. First, the average latency of each cell was determined on the basis of its monaural rate-level functions to allow for subsequent spike-triggered analysis of responses to the HPR stimuli. To this end, spikes were assigned to 50 ms bins of the respective ILD that elicited the spikes (taking into account the latency of the cell). This resulted in a mean ILD-response rate function for each HPR condition. Spike-triggered averages where calculated by selecting all bins with a non zero response and recording the ILD values presented in the nine previous bins plus the bin that showed the non zero response. The average response in each of the ten bins was then used as the Spike-triggered average. The non-spike triggered average was calculated the same way but based on bins that did not show any response. The data are plotted relative to the mean ILD of the last bin in time.

The standard separation D is calculated as previously described (35):

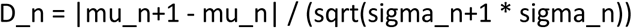

where mu_n+1 and mu_n are the mean values of the responses to two ILD values while sigma_n+1 and sigma_n are their standard deviation. D_n was subsequently smoothed using a 5-sample moving average filter.

The metric D/spike was calculated as:

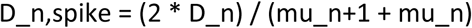

In case of the model, we calculated D based on the assumption of an underlying Poison process where the variance would equal the mean response.

Maximum likelihood estimations (MLE) were used to find the most probable ILD to result in a specific observed response *R*_*obs*_ given all other observed responses R. For this, the joint probability density functions *P*(*R, ILD*) of the observed spike counts R and the presented ILDs was calculated for all responses of one neuron excluding*R*_*obs*_. The ILD that maximizes *P*(*R* = *R*_*obs*_, *ILD*) was than used as the MLE for *R*_*obs*_.

To characterize shifts in ILD functions due to HPR statistics, the threshold ILD, defined as the ILD at which the firing rate differentiates more than 10 % from baseline firing, was determined.

Time courses of adaptation were measured by fitting a single-or double-exponential function to the mean responses rates averaged over all 30 repetitions of a HPR condition. Inbuilt functions in Matlab for the root-mean-square error (rmse) and the adjusted coefficient of determination (R^2^) were used to evaluate the goodness of fits.

### Psychophysical measurements and data analysis

5 normal hearing (within 20dB of ISO/TR 389-5:1998) listeners (2 males and 3 females, mean age 26±4years, right-handed) participated in the measurement of just noticeable ILD differences. The signals consisted of bandpass filtered noise (center frequency 10 kHz, bandwidth 2.208 kHz) which was generated in MatLab (The Mathworks, Inc, Natick, Massachusetts, US) at a sampling rate of 44.1 kHz. The signals were digital to analog converted (Audio 2 Dj, Native Instruments GmbH, Berlin, Germany) before being presented over calibrated circumaural headphones (HDA 200, Sennheiser Electronic GmbH & Co. KG., Wedemark, Germany). The signals where presented at 60 dB SPL average diotic sound pressure level and ILDs were introduced by symmetrically amplifying and attenuating the right and left ear signals by half the desired ILD. Within the experiment, a 2 s long adapter stimulus was followed, after 350ms, by two 50ms probe stimuli which were separated by 100ms. Similar to the physiological experiments, the adapter consisted of concatenated diotic noise busts, each 50 ms in duration, with ILDs that were randomly drawn from one of the two non-uniform HPR distributions. ILD JNDs were determined at two reference ILDs (i.e −20dB ILD and +20dB ILD, see Fig. 2d and e). One of the two probe stimuli was randomly presented at one of the two reference ILDs while the other probe stimulus was systematically varied using a transformed up-down procedure, following a one-up three-down rule, as implemented by the MatLab AFC package(54). To determine the JND, listeners were asked to specify the perceived direction of the probe pair sounds, which allows deducing which of the two probe stimuli was perceived more lateralized. Following the subject’s answer, the variable probe ILD was adjusted until reaching the termination criterion (6 reversals) of the one-up three-down rule. ILD JNDs for each listener, each probe position and each listening condition (i.e. HPR) were calculated as median over six sessions (each session consisting of 3 measurements). For each subject, the effect of listening condition was expressed as normalized change in ILD JND due to co-location of probe position and preceding HPR.

### LSO model

The LSO was modeled using a phenomenological rate model similar to the one used by Wen and colleagues (17) to model adaptation in the ANF (see Fig. S1). The LSO is implemented as a subtraction stage with inputs from the ipsi-and contra lateral ANFs and a sigmoidal activation function (CN and MNTB were omitted to minimize model complexity). The firing rate *R*_*LSO*_(*t*) in spikes per second (sps) of the LSO is calculated as following:

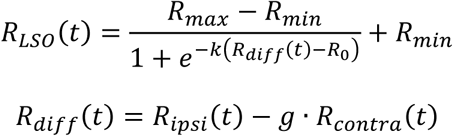

Where *R*_*max*_ and *R*_*min*_ are the maximum and the minimum firing rates, *R*_*0*_ is the rate at zero input and k is the steepness of the sigmoid. *R*_*ipsi*_(*t*) and *R*_*contra*_(*t*) are the firing rates from the ipsi- and the contra-lateral ANFs and *g* is a gain factor to weight the relative strength of the excitatory and inhibitory input. The ANF inputs where each calculated using a dual adaptation model (17), which was fitted to the data shown in figure 2 of (16). As we only fitted the response of one ANF, we switched the saturating nonlinearity used in the original model with a simple logistic function. The parameters for the LSO model where determined by calculating the ILD-rate function of the model and fitting it to the ILD-rate function given by figure 4 in (55) (resulting parameters: 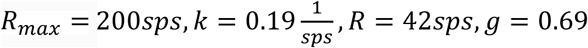). The slow LSO adaptation was implemented to resemble the second adaptation stage of (17). The adaptation parameters where adjusted so that the time course of adaptation was in agreement with the recorded data from the LSO. This resulted in an adaptation time constantan of 4ms, *g*_1_ = 0.3, *g*_2_ = 0.01 with an adaptation threshold of 35sps (see (17) for details on the implementation). Running this model often resulted in zero firing rates for larger negative ILD values which lead to undefined D/spike values so that we introduced a minimal LSO firing rate of *R*_*min*_ = 30*sps*.

### LSO intrinsic metabolic imaging

Changes in NADH levels in the LSO were monitored by imaging of NADH autofluorescence in acute brainstem slices as recently described(41,56). The animals were anesthetized with isoflurane and decapitated. We removed the brains and cut 250-μm thick transverse slices (VT1200S Vibratome, Leica Microsystems GmbH, Wetzlar, Germany). The slices were superfused at room temperature (22– 25 °C) in the recording chamber with gassed (95% O2 and 5% CO2) artificial cerebrospinal fluid (ACSF) solution containing (in mm): 23 sucrose, 125 NaCl, 25 NaHCO3, 2.5 KCl, 1.25 NaH2PO4, 1 MgCl2, 2 CaCl2, 2 glucose (Sigma-Aldrich). Nicotinamide adenine dinucleotide (NADH) was excited with a 365 nm LED and fluorescence images (emission filter: 447±30 nm) were recorded at 2 Hz (pco.edge 5.5; PCO AG, Kehlheim, Germany). LSO neurons were electrically excited by a 20-s stimulation train at 200 Hz with biphasic pulses of 1 ms duration and 5 V amplitude using a bipolar Tungsten electrode placed in the fiber tract targeting the LSO. NADH fluorescence intensity was measured in individual rectangular regions of interest (ROIs), corrected for photobleaching and presented as ΔF/F0 (F0 = fluorescence level at stimulation onset; ΔF = change in fluorescence level relative F0). The occurrence of double minima was automatically detected based on their amplitudes (> 0.05-0.10 %), the time difference between them (> 6-10 s) and the time differences between the minima and the interjacent maximum (> 1-2 s). These parameters were individually adjusted for each slice by analyzing a measurement with larger ROIs and by comparing the automated results with those of visual inspection of the individual traces. Specific blockade of GABA-B receptors was performed by wash in of 10 µM CGP 55845 hydrochlorid [(2*S*)-3-[[(1*S*)-1-(3,4-Dichlorophenyl)ethyl]amino-2-hydroxypropyl](phenylmethyl)phosphinic acid hydrochloride] (Tocris Bioscience) for 20 minutes.

## Acknowledgements

This work was supported by the German research Council (DFG) to B.G. (CRC 870/B02), M.P. (CRC 870/B02 and SPP1608) and J.E. (HE 6731/1-2), the Federal Ministry of Education and Research (BMBF) to B.G. (IFB-LMU, TRFII-18) and the Bavarian Academy of Sciences (to M.P.). We thank C. Leibold for discussions and N. Lesica for comments on earlier versions of the manuscript.

## Author contributions

H.G. performed electrophysiological experiments and analysis. J.E. performed data analysis and modeling. T.R.J. performed stimulus programming and analysis. A.L. designed, conducted and analyzed psychophysical experiments. S.B. performed LSO intrinsic imaging experiments and analyzed the data. L.K. analyzed intrinsic imaging data. B.G. designed and supervised the study. M.P. designed and supervised the study, analyzed the data and wrote the original version of the manuscript.

## Supporting figure captions

**Figure S1.**
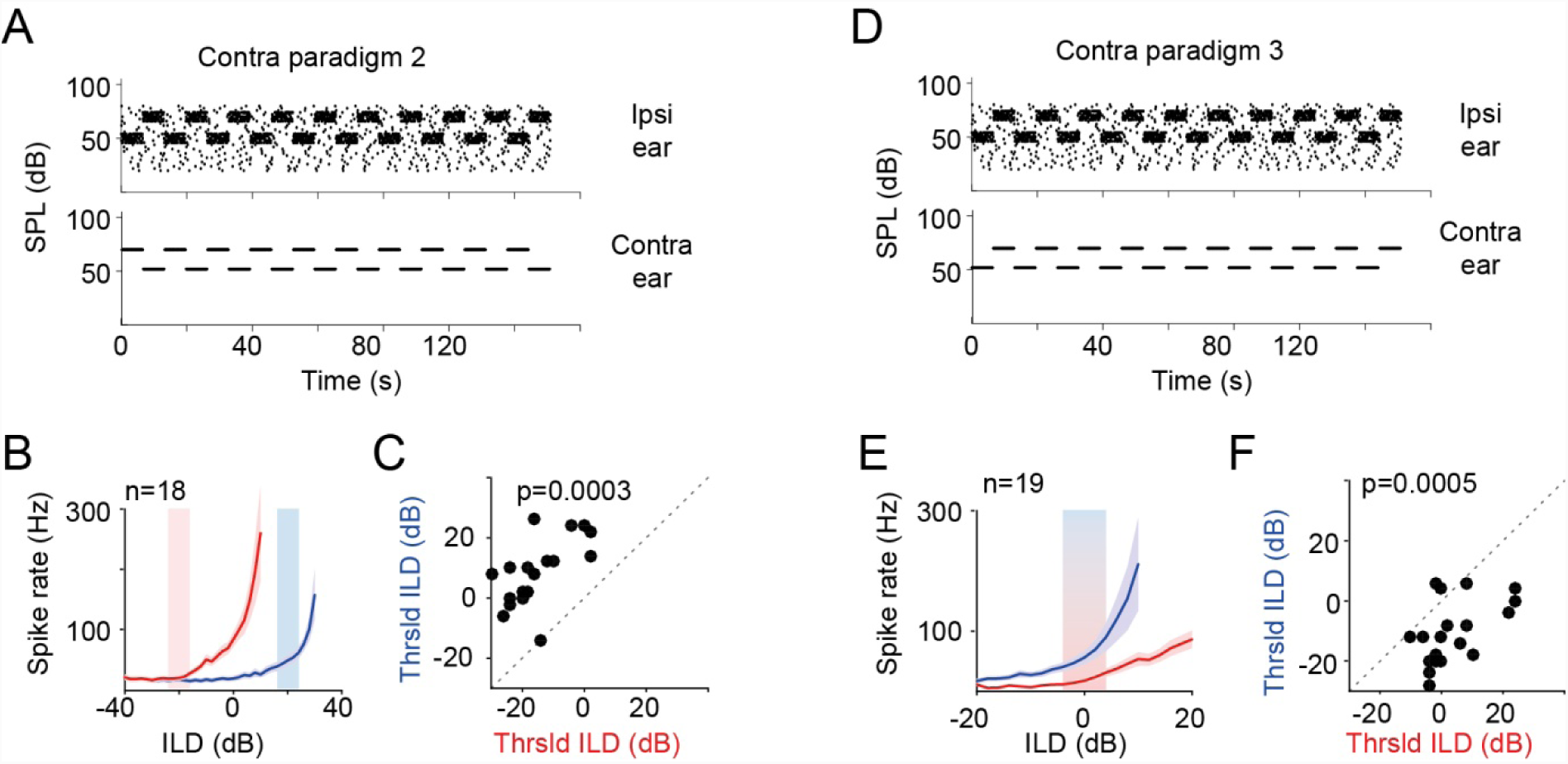
DRA in LSO is specific for binaural statistics. A) and D) Schematics of additional HPR stimulus paradigms. In both, the intensity distribution on the ipsilateral ear was identical to the main HPR paradigm (Fig. 1). However, the intensity distribution on the contralateral ear was only switched between 70dB and 50dB. These intensities were either presented out of phase with the mean HPR intensity on the ipsilateral ear (paradigm 2, panel A) or in phase (paradigm 3, panel D). B) Mean ILD-response functions of all LSO neurons tested with paradigm 2 (n=18 neurons), exhibiting strong DRA as a function of the binaural statistics. Conventions as in figure 2C. C) Scatter plot of threshold ILDs illustrates significant changes with HPR condition in paradigm 2 (p=0.0003, N=18 neurons, paired Wilcoxon signed rank test). Conventions as in figure 1. E) Mean ILD-response functions of all LSO neurons tested with paradigm 3 (n=19 neurons). Conventions as in figure 2C. G) Scatter plot of threshold ILDs illustrates significant changes with HPR condition in paradigm 3 (p=0.0005, N=19 neurons, paired Wilcoxon signed rank test). Conventions as in figure 1.

**Figure S2.**
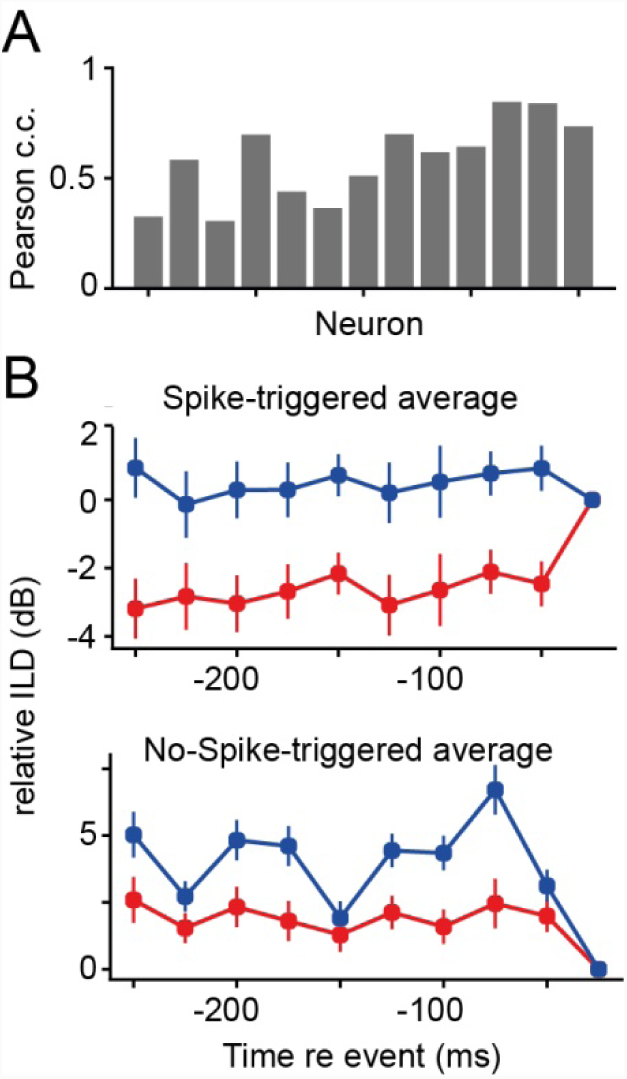
No systematic relationship between ILD sequences and LSO response variability during complex stimulation. A) Histogram of Pearson correlations of LSO responses to the full HPR stimulus. The average correlation of spike counts across three repetitions of the entire stimulus set (20 19 switches) was surprisingly low for the majority of neurons (median Pearson correlation coefficient= 0.62, IQR: 0.26). A spike-triggered analysis of the responses of all neurons established that the likelihood of spiking to any ILD was not systematically associated with a prior occurrence of specific relative ILDs (upper panel, color code represents HPR conditions). Performing the same analysis but triggered by non-spiking to a represented ILD (lower panel) exposed a tendency of non-responsiveness due to presentation of a more positive ILD shortly before.

**Figure S3.**
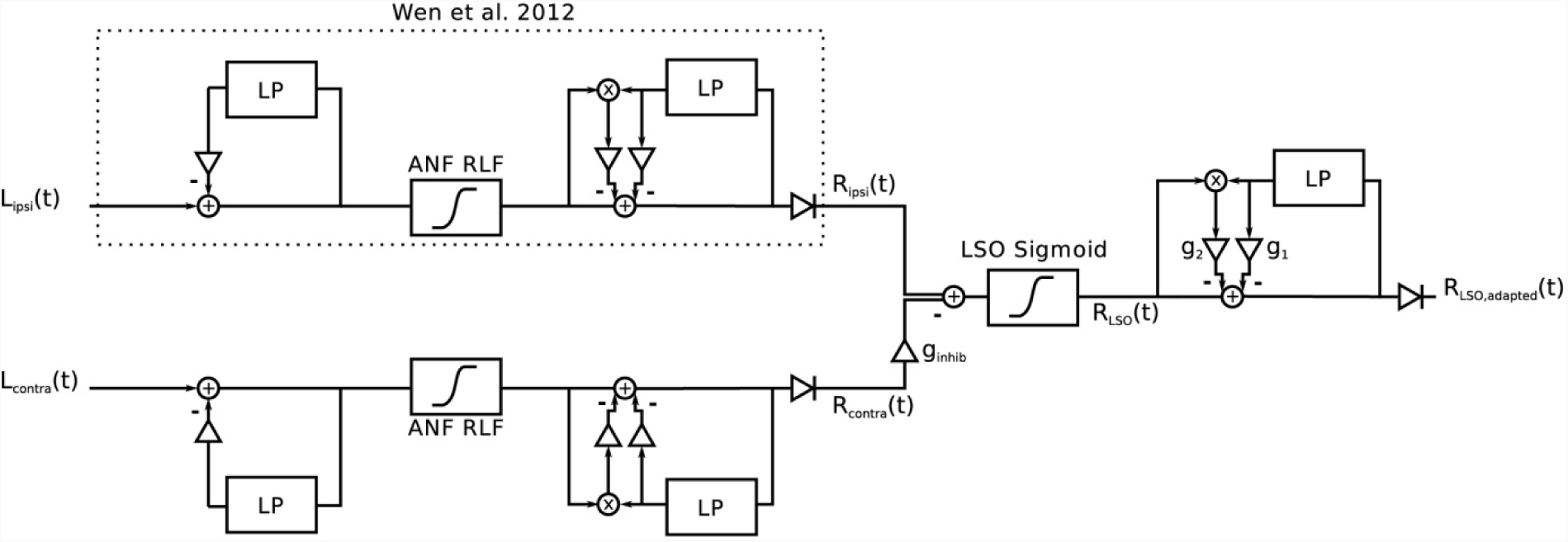
A simple model of the LSO based on existing models of DRA that includes both threshold and gain adaptation. Block diagram of the LSO rate model. The input to the model is given as a sequence of sound levels on the ipsi-(L_ipsi) and contra-lateral (L_contra) ear. A dual adaptation model is used to calculate the auditory nerve fiber (ANF) firing rates R_ipsi and R_contra. The LSO model is implemented as a subtraction stage with the contralateral input weighted by a gain value and a following Sigmoid to model the activation of the neuron. An optional adaptation stage which resembles the rate adaptation stage in the ANF model is used to account for the slow adaptation component present in the LSO measurements.

